# HIV-1 Vpr-induced DNA damage activates NF-κB through ATM-NEMO independent of cell cycle arrest

**DOI:** 10.1101/2023.05.23.541990

**Authors:** Carina Sandoval, Karly Nisson, Oliver I. Fregoso

**Affiliations:** Molecular Biology Institute, University of California, Los Angeles, California, USA; Department of Microbiology, Immunology, and Molecular Genetics, University of California, Los Angeles, California, USA

## Abstract

Lentiviral accessory genes enhance replication through diverse mechanisms. HIV-1 accessory protein Vpr modulates the host DNA damage response (DDR) at multiple steps through DNA damage, cell cycle arrest, the degradation of host proteins, and both the activation and repression of DDR signaling. Vpr also alters host and viral transcription; however, the connection between Vpr-mediated DDR modulation and transcriptional activation remains unclear. Here, we determined the cellular consequences of Vpr-induced DNA damage using Vpr mutants that allow us to separate the ability of Vpr to induce DNA damage from cell cycle arrest and other DDR phenotypes including host protein degradation and repression of DDR. RNA-sequencing of cells expressing Vpr or Vpr mutants identified that Vpr alters cellular transcription through mechanisms both dependent and independent of cell cycle arrest. In tissue-cultured U2OS cells and primary human monocyte-derived macrophages (MDMs), Vpr-induced DNA damage activates the ATM-NEMO pathway and alters cellular transcription via NF-κB/RelA signaling. HIV-1 infection of primary MDMs validated Vpr-dependent NF-κB transcriptional activation during infection. Both virion delivered and *de novo* expressed Vpr induced DNA damage and activated ATM-NEMO dependent NF-κB transcription, suggesting that engagement of the DDR and transcriptional reprogramming can occur during early and late stages of viral replication. Together, our data identifies a mechanism by which Vpr activates NF-κB through DNA damage and the ATM-NEMO pathway, which occur independent of cell cycle arrest. We propose this is essential to overcoming restrictive environments, such as in macrophages, to enhance viral transcription and replication.

**IMPORTANCE:** The HIV accessory protein Vpr is multi-functional and required for viral replication *in vivo*, yet how Vpr enhances viral replication is unknown. Emerging literature suggests that a conserved function of Vpr is engagement of the host DNA damage response (DDR). For example, Vpr activates DDR signaling, causes DDR-dependent cell cycle arrest, promotes degradation of various DDR proteins, and alters cellular consequences of DDR activation. However, a central understanding of how these phenotypes connect and how they affect HIV-infected cells remains unknown. Here, we found that Vpr-induced DNA damage alters the host transcriptome by activating an essential transcription pathway, NF-κB. This occurs early during infection of primary human immune cells, suggesting NF-κB activation and transcriptome remodeling are important for establishing productive HIV-1 infection. Together, our study provides novel insights into how Vpr alters the host environment through the DDR, and what roles Vpr and the DDR play to enhance HIV replication.

## INTRODUCTION

The DDR is a signaling cascade activated in response to exogenous and endogenous genotoxic stressors, such as DNA breaks. The DDR consists of sensor proteins that sense the damaged DNA, transducer proteins that transmit the signal of damaged DNA, and effector proteins that elicit a cellular response. Typically, the DDR is divided into three main pathways based on the mediator kinases activated: Ataxia telangiectasia and Rad3 related (ATR), ataxia telangiectasia mutated (ATM), and DNA-dependent protein kinase (DNA-PK). Single-strand DNA breaks (SSBs) are primarily sensed by RPA to activate ATR signaling, while double-strand DNA breaks (DSBs) can be sensed by the MRN (MRE11, RAD51, and NBS1) or Ku70/80 complexes to activate either ATM or DNA-PK signaling, respectively (1,2). However, a significant amount of crosstalk exists between these pathways. DDR signaling leads to various cellular responses, such as DNA repair, cell cycle arrest (3), transcriptional changes (4,5), apoptosis, or senescence (6).

Many diverse RNA and DNA viruses have evolved to modulate the DDR to enhance viral replication (7,8). Primate lentiviruses, such as HIV-1, primarily engage the DDR through the accessory protein Vpr (9,10). Vpr is evolutionarily conserved by extant primate lentiviruses and is both delivered by the incoming virion, allowing it to act early in viral replication, and expressed *de novo* from the integrated provirus, allowing it to act later in viral replication (11). Viruses lacking Vpr have no appreciable replication defects in T cells and most transformed cell lines, however Vpr is required for replication *in vivo* (12,13) and viruses lacking Vpr have decreased proviral transcription in monocyte derived macrophages (MDMs) (13,14) and dendritic cells (14). This suggests an important role for Vpr in viral transcription. Moreover, studies of Vpr orthologs and mutants in isolation, whether overexpressed or delivered by virus-like particles (VLPs), identified that Vpr engages the host DDR at multiple, potentially unique, steps.

For example, ATR activation by Vpr leads to cell cycle arrest and requires recruitment of the CRL4A^DCAF1^ complex, which has a primary role in DNA repair (15). CRL4A^DCAF1^ complex recruitment by Vpr also leads to the degradation of many host proteins involved in the DDR (18–27) and is required for Vpr-mediated repression of DSB repair (16). In contrast, we have previously shown that the ability of Vpr to induce DNA damage does not correlate with ATR activation, cell cycle arrest, or repression of DSB repair (16), suggesting that Vpr-induced DNA damage may have unique roles in enhancing lentiviral replication.

Vpr, DNA damage, and proviral transcription are connected through NF-κB in that both Vpr and DNA damage promote transcriptional changes involving NF-κB (17,18), which is an essential transcription factor for HIV-1 proviral transcription(19,20). In response to DSBs, ATM signaling promotes NF-κB transcriptional upregulation through the ATM-NEMO pathway (21). ATM and NEMO interact in the nucleus before translocating to the cytoplasm (22,23) to subsequently activate RelA nuclear translocation, promoter binding, and NF-κB transcriptional activation. Previous literature suggests that HIV-1 Vpr activates and represses NF-κB pathways through phosphorylation and ubiquitylation of TAK1 to enhance NF-κB signaling (24) and altering the availability of the NF-κB p50-RelA heterodimer to inhibit NF-κB signaling (25), respectively. This proposes a testable model where Vpr-induced DNA damage alters the cellular environment to enhance viral replication by altering transcription through ATM-NEMO and NF-κB. Here, we aimed to identify the cellular consequences of Vpr-induced DNA damage and the connection between DNA damage and ATM activation, cell cycle arrest, and transcriptional changes.

To determine the consequences of Vpr-induced DNA damage on cellular transcription, we used previously described Vpr mutants that allow us to uncouple DNA damage from the large-scale cellular changes caused by cell cycle arrest. We found that virion-associated or *de novo* expressed Vpr does not require cell cycle arrest to induce DNA breaks and activate DDR signaling in U2OS tissue-culture cells and primary human MDMs. RNA-sequencing (RNA-seq) identified that wild-type HIV-1 Vpr and Vpr mutants that do not induce cell cycle arrest still alter NF-κB associated cellular transcription. In support of this, we showed that Vpr proteins that induce DNA damage activate RelA nuclear translocation and upregulate NF-κB target genes, such as BIRC3 and CXCL8. We further assessed the requirement for ATM and NEMO signaling in Vpr-mediated NF-κB activation in U2OS cells and primary human MDMs. We found that Vpr-induced DNA damage activates canonical ATM signaling, and that inhibition of NEMO resulted in loss of NF-κB transcriptional upregulation. HIV-1 infection and virus-like particle (VLP) delivery of Vpr in MDMs validated the Vpr-dependent upregulation of NF-κB target genes early during infection. Together, our data support a model where Vpr-induced DNA damage activates NF-κB through the ATM-NEMO pathway, independent of cell cycle arrest and associated phenotypes. This study further informs how lentiviral accessory proteins engage the DDR at multiple and unique steps, and clarifies how this engagement remodels the host environment and immune pathways to promote viral replication.

## RESULTS

### HIV-1 Vpr alters cellular transcription independent of cell cycle arrest

Given the connections between the DDR and cellular transcription, we set out to understand whether the ability of Vpr to induce DNA damage alters cellular transcription, and whether these transcriptional changes are distinct from those caused by Vpr-mediated cell cycle arrest. To do this, we leveraged a subset of previously established HIV-1 Vpr mutants that inhibit Vpr-mediated cell cycle arrest: Q65R, H71R, and S79A (all generated in the transmitted founder HIV-1 Q23-17 background (16)). These mutants have often been used interchangeably throughout the Vpr literature; however, they have not been fully characterized in the same system. We therefore first evaluated the ability of HIV-1 Q23-17 Vpr wild type (WT) and mutants to induce DNA damage, activate DDR signaling, cause cell cycle arrest, and interact with DCAF1 (through which Vpr binds the CRL4A^DCAF1^ E3 ubiquitin ligase complex (26)), as well as their subcellular localization. U2OS cells were infected with a recombinant adeno-associated virus (rAAV) expressing either HIV-1 Q23-17 Vpr WT or mutants(10,16) (Fig. S1A). We confirmed that all mutants lost the ability to cause cell cycle arrest (Fig. S1B), as has been previously described(16,27–29). DNA damage was assessed by the comet assay while DDR activation was assessed by immunofluorescence 24 hours post infection. Consistent with our previous results (16), HIV-1 Vpr WT and S79A induce DNA breaks (Fig. 1A) and activate the DDR marker γH2A.x (Fig. 1B), while Q65R does not. In addition, we found that H71R induces DNA breaks (Fig. 1A) and activates γH2A.x (Fig. 1B) at levels similar to Vpr WT. Only the H71R mutant is proposed to lose interaction with the DCAF1 component of the CRL4A^DCAF1^ E3 ubiquitin ligase complex and thus lose the ability to cause cell cycle arrest and degrade host proteins (16,30–32). In our hands, HIV-1 Q23-17 Vpr WT and S79A recruit similar amounts of DCAF1, H71R recruits a decreased amount of DCAF1 when compared to Vpr WT and S79A, and Q65R recruits background levels of DCAF1 (Fig. S1C). Finally, we found that all Vpr mutants except for Vpr Q65R show similar subcellular localization as WT Vpr (Fig. S1D). Together, our data indicate that, when compared to WT Vpr, Vpr S79A can be used to assess the aspects of Vpr phenotypes that are specific to cell cycle arrest, Vpr H71R can be used to assess those aspects specific to cell cycle arrest and diminished DCAF1 recruitment, and Vpr Q65R can be used as a functionally dead control as it has lost all Vpr-associated phenotypes (Table 1).

**Fig 1:**
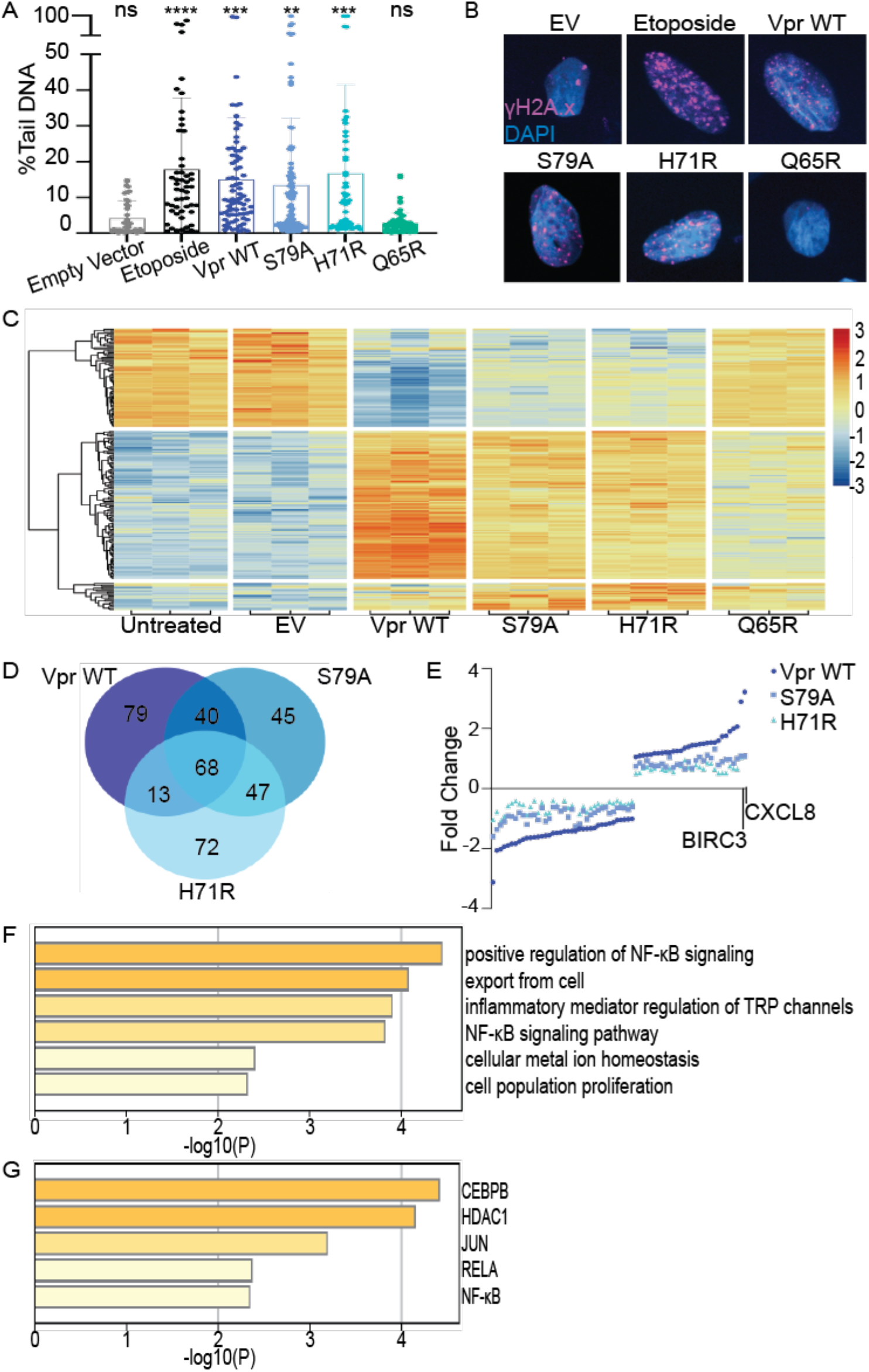
HIV-1 Vpr induces DNA damage and alters cellular transcription independent of cell cycle arrest. (A) Comet assay of U2OS cells infected with rAAV expressing 3xFLAG-tagged Vpr WT, S79A, H71R, Q65R, empty vector (EV) (negative control), or 50uM etoposide (positive control). Percent tail DNA was quantified at 24 hours post infection (hpi) using the OpenComet plug-in for the ImageJ software. Each circle represents one cell. N=3, one representative experiment shown. (B) Representative immunofluorescence images of U2OS cells infected under the same conditions as Fig. 1A; γH2A.x (magenta) and nuclei stained with DAPI (blue). Images were taken at 63x magnification. N=3, one representative experiment shown. (C) RNA-seq of U2OS cells at 36 hpi. Heat map displays Log_2_ fold changes of upregulated genes in red and downregulated genes in blue. Each column represents a biological replicate. (D) Venn diagram of the top 100 upregulated and top 100 downregulated genes among Vpr WT, S79A, H71R, and Q65R. (E) Dot plot of the 68 conserved differentially expressed genes among Vpr WT, S79A, and H71R. BIRC3 and CXCL8 are highlighted as the two most upregulated genes (Log_2_ fold change) under all conditions. (F) Gene ontology and (G) TRRUST analysis of the 30 upregulated genes performed with Metascape software. Asterisk indicate statistical significance compared to EV control, as determined by one-way ANOVA test (NS, nonsignificant; ** P< 0.001, *** P< 0.0004, **** P< 0.0001). Related to Figures S1 and S2.

**Table 1.** HIV-1 Vpr WT and mutant phenotypes. (+) indicates a phenotype conferred by Vpr, (-) indicates a phenotype not conferred by Vpr, and (-/+) indicates an intermediate Vpr phenotype.

|  | Localization as<br>WT | Cell Cycle<br>Arrest | Engages DCAF1 | DNA<br>Damage | NF-κB<br>activation |
| --- | --- | --- | --- | --- | --- |
| HIV-1 Vpr WT | + | + | + | + | + |
| S79A | + | - | + | + | + |
| H71R | + | - | - / + | + | + |
| Q65R | - | - | - | - | - |

We next asked if Vpr mutants that induce DNA damage in the absence of cell cycle arrest (H71R and S79A) alter the cellular transcriptome. U2OS cells were infected with rAAV expressing HIV-1 Vpr WT, H71R, S79A, or Q65R, and RNA was collected at 12, 24, and 36 hours post infection for analysis by bulk RNA-sequencing (RNA-seq). Vpr WT, H71R, and S79A significantly altered cellular transcription 36 hours post infection when compared to empty vector or untreated cells, while Q65R was indistinguishable from empty vector or untreated cells (Fig. 1C-E, Fig. S2A-B, and Supplemental Files S1 & S2). The largest effect on cellular transcription was seen with Vpr WT (950 differentially expressed genes, Log_2_(2) fold change p-value < 0.004 and FDR < 0.01), while H71R and S79A showed an intermediate effect (361 differentially expressed genes, Log_2_(1.35) p-value < 0.0007 and FDR < 0.01 and 233 differentially expressed genes, Log_2_(1.5) p-value < 0.0008 and FDR < 0.006), respectively. All cutoffs failed to identify significant differentially expressed genes in cells expressing Vpr Q65R, empty vector or untreated cells. These data suggest that Vpr the alters cellular transcriptome via mechanisms that are dependent and independent of cell cycle arrest.

We focused on genes differentially expressed among Vpr WT, H71R, and S79A, as these genes are indicative of changes in cellular transcription that occur independent of cell cycle arrest and are potentially altered by Vpr-induced DNA damage. Out of the top 200 differentially expressed genes for each condition, 68 genes with p-value < 0.01 and FDR < 4.50E-05 were shared among Vpr WT, H71R, and S79A, while 100 additional genes were shared by two out of three conditions (Fig. 1D and 1E, Supplemental Files S1 & S2). Gene ontology (Fig. 1F) and Transcriptional Regulatory Relationships Unraveled by Sentence-based Text-mining (TRRUST)(33) (Fig. 1G) analyses of shared upregulated genes identified that Vpr WT, S79A, and H71R mutants were significantly enriched for positive regulation of RelA/NF-κB signaling and NF-κB signaling pathways (Fig. 1F-G, Fig. S2 A-B and Supplemental Files S1 & S2). TRRUST analysis further identified upregulation of transcription factors that bind and activate the HIV-1 LTR, such as CEBPB (34), SP1 (35), and JUN (36), as well as HDAC1, which modifies the HIV-1 LTR directly (37) (Fig. 1G). Moreover, Protein-protein Interaction Enrichment Analysis of the shared differentially expressed genes identified that Vpr downregulates expression of histones belonging to the H2A, H2B, H4, and H1 family that are involved in nucleosome assembly and DNA Damage/Telomere Stress Induced Senescence (Fig. S2C); consistent with previous literature that Vpr modulates components of the chromatin environment (Romani, Johnson). Collectively, our data show that Vpr alters cellular transcription in the absence of cell cycle arrest and suggest that Vpr-induced DNA damage robustly activates RelA/NF-κB-mediated signaling.

### HIV-1 Vpr-induced DNA damage activates RelA and promotes NF-κB transcription

Given NF-κB signaling was the strongest upregulated pathway by Vpr WT, H71R, and S79A mutants, that NF-κB signaling is activated by DNA damage, and the precedence for Vpr-mediated modulation of NF-κB (24), we next directly assayed whether Vpr activates RelA/NF-κB in the absence of cell cycle arrest. We focused on H71R because it has lost the ability to cause cell cycle arrest and has diminished DCAF1 binding and host protein degradation (30–32), but maintains the ability to induce DNA damage and activate DDR signaling. NF-κB activation was first validated by qRT-PCR for two NF-κB target genes identified in our RNA-seq, BIRC3 and CXCL8 (Fig. 1E), which play important roles in innate immunity and cell survival (38,39). U2OS cells were infected with rAAV expressing Vpr WT, H71R, and Q65R for 24 or 36 hours. Consistent with the RNA-seq, Vpr WT and H71R upregulate BIRC3 and CXCL8 compared to untreated cells or Q65R mutant at 36hpi (Fig. 2A). To directly assess NF-κB activation, we assayed for RelA localization by immunofluorescence. As RelA is cytoplasmic when inactive and nuclear when active, we expect that Vpr WT and H71R will lead to nuclear translocation of RelA. Indeed, we found that Vpr WT and H71R activate nuclear translocation of RelA similar to the positive control, etoposide, while Vpr Q65R and empty vector do not (Fig 2B). These data demonstrate that Vpr activates RelA and NF-κB transcription in the absence of cell cycle arrest, suggesting that Vpr-induced DNA damage is sufficient to drive NF-κB activation.

**Fig 2:**
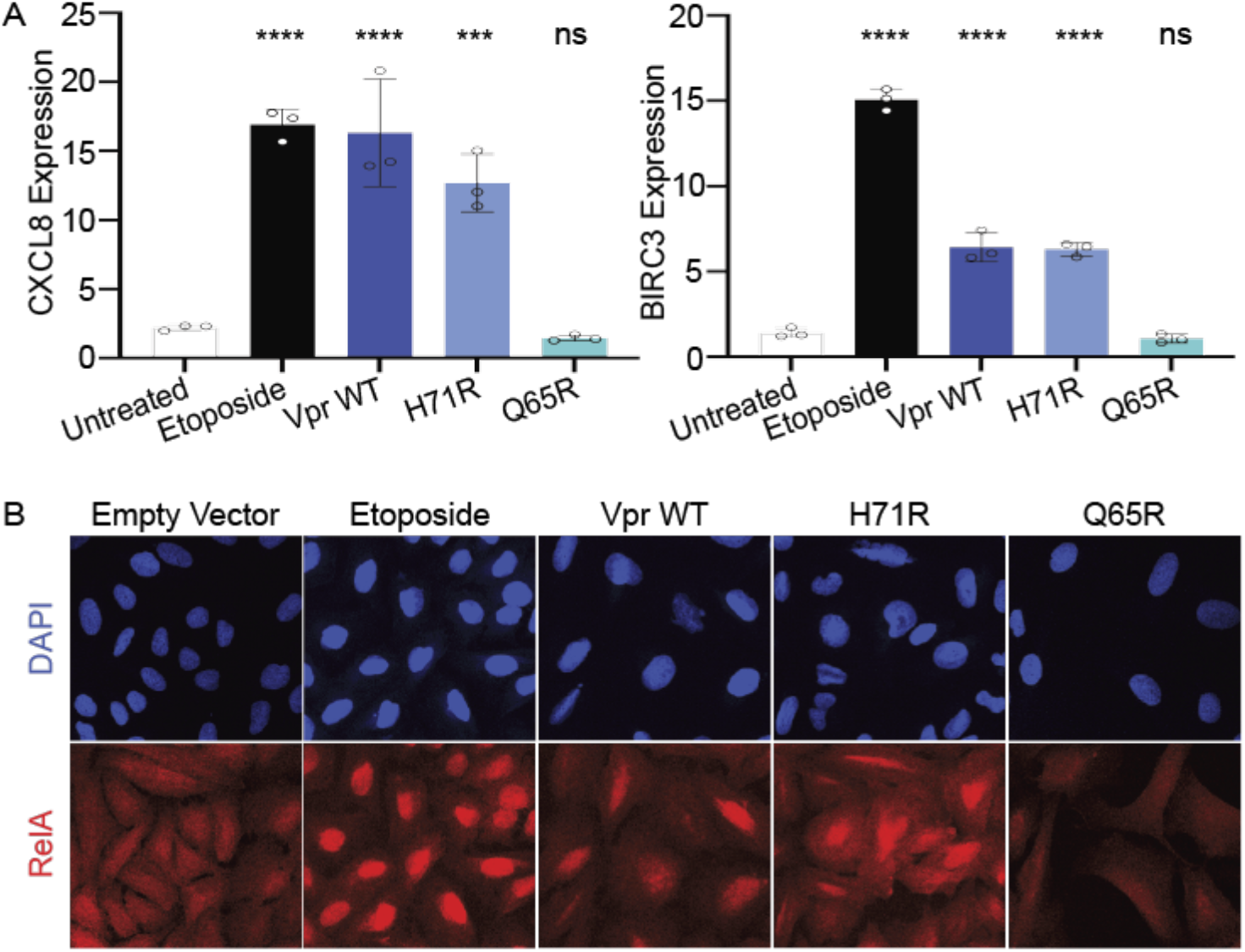
HIV-1 Vpr activates RelA and promotes NF-κB transcription independent of cell cycle arrest. (A) qRT-PCR validation of upregulated NF-κB target genes BIRC3 and CXCL8. Normalized expression (ΔΔCt) was calculated by normalizing to GAPDH followed by calculating fold change to untreated or empty vector treated cells. Cells treated under the same conditions as Fig. 1C. N=3, one representative experiment shown with technical triplicates. (B) Representative immunofluorescence images of U2OS cells infected under the same conditions as Fig. 1A; RelA (red) and nuclei stained with DAPI (blue). Images were taken at 63x magnification. N=3, one representative experiment shown. Asterisk indicate statistical significance compared to untreated control, as determined by one-way ANOVA test (NS, nonsignificant; *** P< 0.0003, **** P< 0.0001).

### HIV-1 Vpr-induced DNA damage activates ATM-NEMO signaling

NF-κB is activated in response to many pathways and signals involved in DNA repair and innate immunity (40). One such pathway is ATM-NEMO, where ATM stimulates NEMO to activate RelA and thus NF-κB signaling (23). Previous work has shown that Vpr activates markers of both ATR and ATM signaling (16,41). While ATR is required for Vpr-induced cell cycle arrest, the extent to which Vpr activates ATM and the cellular consequences of ATM activation without cell cycle arrest are unclear. We hypothesized that Vpr-induced DNA damage activates the ATM-NEMO pathway, which consequently promotes RelA/NF-κB transcription.

To determine if Vpr-induced DNA damage activates ATM signaling in the absence of cell cycle arrest, we measured activation and colocalization of the DNA damage sensors NBS1 and MRE11 and the downstream signaling transducers, 53BP1 and γH2A.x. U2OS cells stably expressing NBS1-GFP or 53BP1-GFP (42,43) were infected with rAAV expressing Vpr WT, H71R, and Q65R and DDR activation was assessed through live-cell imaging over 56 hours. We found that Vpr WT and H71R promote the formation of NBS1 and 53PB1 foci similar to the positive control etoposide, while the non-functional Q65R Vpr mutant was indistinguishable from untreated control cells (Fig. 3A and Fig. S3A-C). Moreover, both damage sensors, MRE11 and NBS1, and transducers, 53PB1 and γH2A.x, colocalize in Vpr WT and H71R similar to etoposide treated cells (Fig. 3B&C). This suggests that Vpr-induced DNA damage activates, but does not dysregulate, classical ATM signaling.

**Fig 3:**
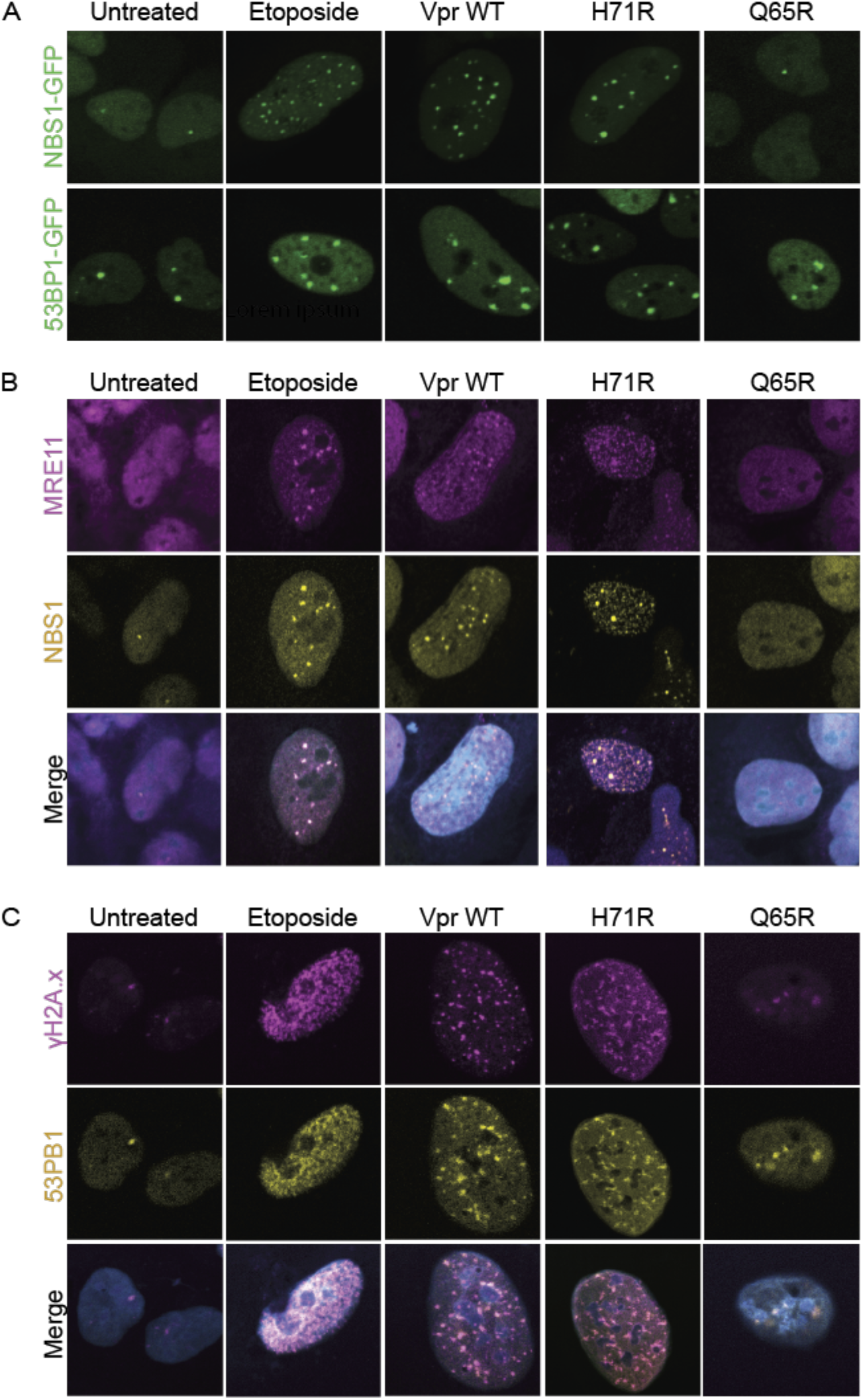
Vpr-induced DNA damage activates ATM signaling independent of cell cycle arrest. (A) Live-cell imaging of U2OS cells stably expressing NBS1-GFP or 53PB1-GFP. Cells were infected under the same conditions as Fig. 1A and representative images were taken at 33 hpi at 63x magnification. N=3, one representative experiment shown. (B) Representative images of co-localization of DNA damage sensors MRE11 (magenta) and NBS1 (yellow) in U2OS NBS1-GFP or (C) DNA damage transducers γH2A.x (magenta) and 53PB1 (yellow) in U2OS 53PB1-GFP cells infected under the same conditions as Fig. 1A. Images taken at 24 hpi at 63x magnification. N=3, one representative experiment shown. Related to Figure S3.

To determine if NEMO is required for Vpr to upregulate NF-κB transcription, we asked whether Vpr could upregulate NF-κB target genes BIRC3 and CXCL8 when NEMO was absent or inhibited. U2OS parental and NEMO knockout cells (44) were infected with rAAV expressing Vpr WT and mutants (Fig. S4A). In the absence of NEMO, Vpr WT and H71R, but not Q65R, still activated the DDR, indicating NEMO is not required for Vpr-induced DDR signaling (Fig. 4A and Fig. S4B). As expected, NEMO knockout inhibited TNFα and etoposide-induced TNFα expression, confirming loss of NEMO function (Fig. 4B). In the absence of NEMO, Vpr WT and H71R lost the ability to upregulate BIRC3 and CXCL8 (Fig 4C), suggesting that NEMO is required for Vpr-induced NF-κB signaling. To further confirm this observation, we inhibited NEMO using a cell-permeable NEMO binding domain (NBD) inhibitor peptide (1). Similar to NEMO knockout cells, NBD peptide, but not a cell permeable peptide negative control, inhibited TNFα expression following TNFα or etoposide treatment (Fig. 4D). To assess whether NBD peptide inhibited Vpr-induced NF-κB activation, U2OS cells were pretreated with NBD peptide and infected with rAAV expressing Vpr WT and mutants. Consistent with NEMO knockout, the NBD peptide inhibited upregulation of BIRC3 and CXCL8 by Vpr WT and H71R (Fig. 4E), and this inhibition was indistinguishable from loss of NEMO (Fig. S4C). Together, this data suggests that Vpr requires NEMO activation to upregulate transcription of NF-κB target genes.

**Fig 4:**
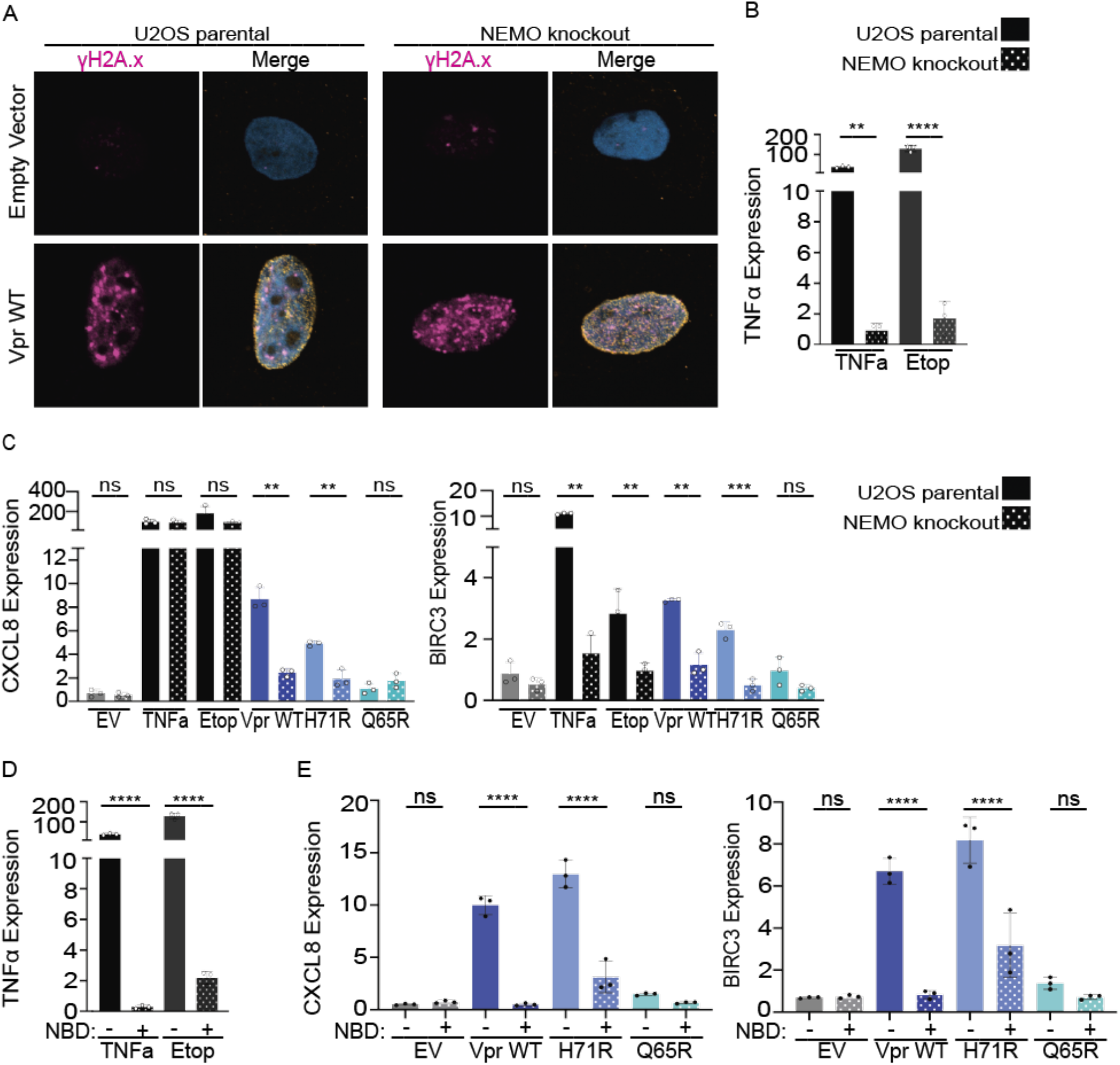
Vpr upregulates NF-κB transcription dependent on NEMO, yet independent of cell cycle arrest. (A) Representative images of U2OS parental or NEMO knockout cells infected with EV or Vpr WT displaying γH2A.x (magenta), Vpr FLAG (yellow), and DAPI (blue). Images taken at 24hpi at 63x magnification. N=3, one representative experiment shown. (B) qRT-PCR of TNFα in U2OS parental or U2OS NEMO knockout cells treated with TNFα or Etoposide (positive controls). RNA was isolated at 36hpi. Asterisk indicate statistical significance compared to U2OS parental cells. (C) qRT-PCR of BIRC3 and CXCL8 in U2OS parental or NEMO knockout cells. Cells were treated under the same conditions as Fig. 1C. N=3, one representative experiment shown with technical triplicates. Asterisk indicate statistical significance compared to U2OS parental cells. (D) qRT-PCR of TNFα in the presence of Nemo-Binding inhibitor peptide (NBD, +) or the NBD negative control peptide (-) at 36 hpi. U2OS cells were pretreated with 50µM of NBD peptide or NBD negative control peptide for 2 hours prior treatment with TNFα or etoposide. Asterisk indicate statistical significance compared to U2OS parental cells. (E) qRT-PCR of BIRC3 and CXCL8 in the presence of NBD inhibitor peptide (+) or the NBD negative control peptide (-) at 36 hpi. U2OS cells were pretreated as in Fig. 4D and infected with rAAV expressing 3xFLAG-tagged Vpr WT, H71R, Q65R, EV (negative control). Normalized expression to GAPDH. N=3, one representative experiment shown with technical triplicates. Asterisk indicate statistical significance compared to NBD negative control, as determined by one-way ANOVA test (NS, nonsignificant; ** P< 0.001, *** P< 0.0003, **** P< 0.0001). Related to Figure S4.

### Virion delivered HIV-1 Vpr activates ATM-NEMO NF-κB signaling in primary MDMs

Together our data proposes a model where Vpr-induced DNA damage activates ATM and NEMO signaling, resulting in RelA nuclear translocation and NF-κB transcriptional upregulation. We next tested this model in primary human MDMs (Fig. 5A), where Vpr enhances viral transcription and replication (46). Primary human MDMs (Fig. S5A) were infected with virus-like particles (VLPs) carrying physiological levels of Vpr WT, H71R, or Q65R (Fig. 5B), and assayed for DNA damage, NF-κB activation, and NEMO dependence (Fig. 5A). Similar to overexpression of Vpr in U2OS tissue culture cell lines, VLP-delivered Vpr WT and H71R mutant, but not Q65R or empty VLPs, induced DNA breaks and activated γH2A.x in primary human MDMs (Fig. 5C-D). Consistent with our data in U2OS cells, VLP-delivered Vpr WT and H71R, but not Q65R or empty VLPs, upregulated BIRC3 and CXCL8 similar to the TNFα positive control in primary human MDMs (Fig. 5E and Fig. S5B). Finally, NBD peptide delivery prior to VLP infection blocked Vpr WT and H71R-mediated BIRC3 and CXCL8 transcriptional activation, but not Vpr-induced DNA damage in primary human MDMs (Fig. 5D-E and Fig. S5B). These data indicate that physiological levels of virion-delivered Vpr in primary human MDMs induce DNA damage that activates NF-κB transcription in a NEMO dependent manner.

**Fig 5:**
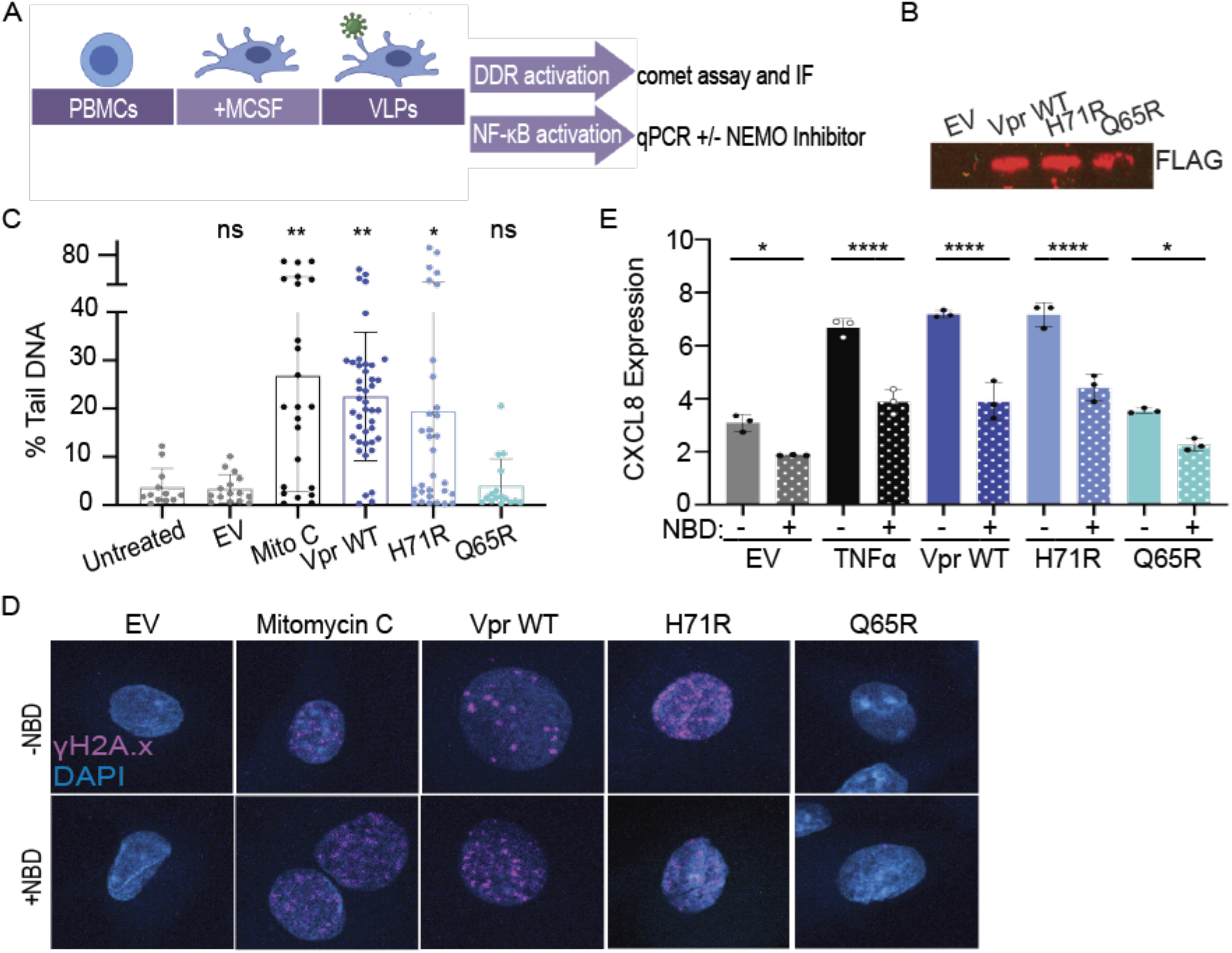
Vpr-induced DNA damage activates ATM-NEMO signaling and NF-κB transcription in primary human MDMs early during infection. (A) Experimental schematic depicting the isolation and differentiation of primary human monocyte-derived macrophages (MDMs) from peripheral blood mononuclear cells (PBMCs) followed by delivery of Vpr via Virus-like Particles (VLPs). Infected MDMs were assayed for induction of DNA damage, activation of DDR signaling, and the upregulation of NF-κB target genes in the context of NEMO-Binding inhibitor peptide (NBD) or the NBD negative control peptide. (B) Western blot of 3X FLAG-Vpr WT and mutants packaged in VLPs. (C) Comet assay of MDMs treated with VLPs packaging 3xFLAG-tagged Vpr WT, H71R, Q65R, empty VLPs (negative control), or 25uM Mitomycin C (Mito C, positive control) for 8 hours. Comet assay analysis was performed as in Fig. 1A. N=2, one representative experiment shown. (D) Representative immunofluorescence images of MDMs infected with VLPs as in Fig. 5B and treated with the NBD inhibitor peptide (+) or the NBD negative control peptide (-). γH2A.x (magenta) and nuclei stained with DAPI (blue). Images were taken at 100x magnification. N=3, one representative experiment shown. (E) qRT-PCR for CXCL8 of MDMs treated with NBD peptide as in Fig. 5C with TNFα positive control. N=3, one representative experiment shown with technical triplicates. Normalized expression to GAPDH. Asterisk indicate statistical significance compared to NBD negative control, as determined by one-way ANOVA test (NS, nonsignificant; * P<0.03, ** P < 0.005, **** P< 0.0001). Related to Figure S5.

To determine if HIV-1 infection activates NF-κB target genes, we infected primary human MDMs from four different donors with HIV-1 ΔEnv and assayed for transcriptional changes at early (8 hours), intermediate (16 hours), and late (24 and 48 hours) timepoints post infection. HIV-1 infection was monitored by flow cytometry for HIV-1 core proteins and qRT-PCR for unspliced HIV-1 genomic transcripts (Fig. S6A-B), while NF-κB activation was measured by qRT-PCR for CXCL8 and BIRC3. HIV-1 ΔEnv infection resulted in upregulation of CXCL8 and BIRC3 in MDMs as early as 8hpi in all donors, despite some donor-to-donor variability in the magnitude of upregulation (Fig. 6A and Fig. S6C), suggesting that HIV-1 infection upregulates NF-κB target genes early during infection, consistent with VLP delivery of Vpr.

**Fig 6:**
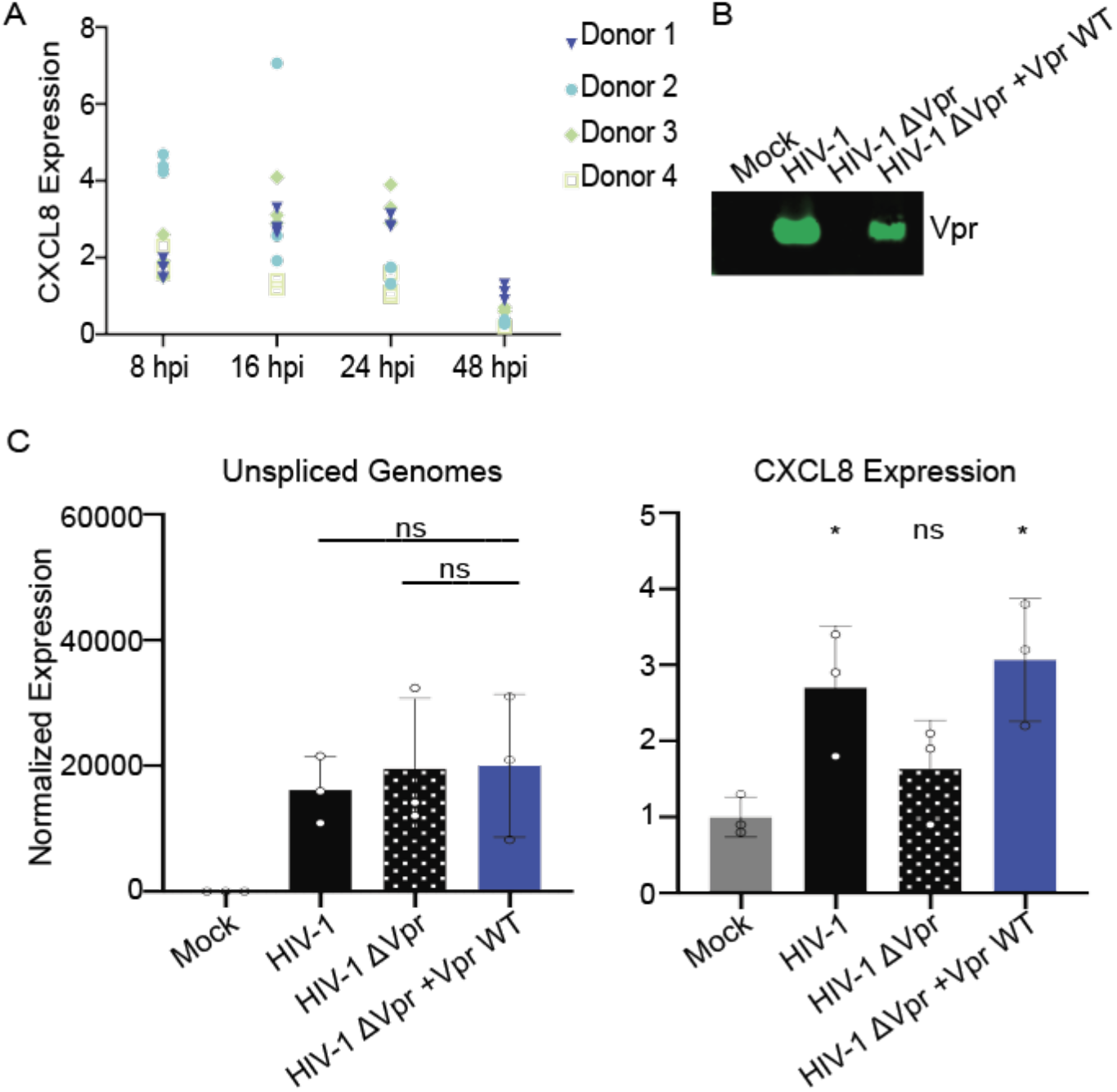
Vpr is necessary and sufficient for activation of NF-kB target genes by HIV-1 in primary human MDMs. (A) qRT-PCR for CXCL8 of MDMs infected with HIV-1 ΔEnv for 8, 16, 24, and 48 hpi from four separate donors. Normalized expression to GAPDH. (B) Western blot of Vpr packaged in mock, HIV-1, HIV-1 ΔVpr, HIV-1 ΔVpr +Vpr WT viruses. (C) qRT-PCR for unspliced HIV-1 genomic transcripts and CXCL8 from MDMs infected with MOI 5U/mL (RT activity) of HIV-1, HIV-1 ΔVpr, HIV-1 ΔVpr +Vpr WT viruses or mock (negative control). The mean (N=3) of three separate donors is shown. Normalized expression to GAPDH. Asterisk indicate statistical significance compared to mock negative control, as determined by one-way ANOVA (NS, nonsignificant; * P < 0.005). Related to Figure S6.

To test whether Vpr was responsible for the upregulation of NF-κB target genes early during HIV-1 ΔENV infection, we generated two additional viruses: HIV-1 ΔVpr and HIV-1 ΔVpr + Vpr packaged in trans (Fig. 6B). We infected primary human MDMs from three additional donors with either HIV-1 ΔENV, HIV-1 ΔENV ΔVpr, HIV-1 ΔENV ΔVpr + Vpr, or mock and assayed for infection and NF-κB activation at 8 hpi. All three viruses infected cells similarly at 8hpi as shown by qRT-PCR for unspliced HIV-1 genomic transcripts (Fig. 6C) and by flow cytometry for HIV-1 core proteins at 48hpi (Fig. S6D). HIV-1 infection upregulated CXCL8 and BIRC3 gene expression, while HIV-1 ΔVpr did not (Fig. 6C and Fig S6E-F). Moreover, upregulation of CXCL8 and BIRC3 were rescued by infection of HIV-1 ΔVpr + Vpr (Fig. 6C). These data suggest that Vpr is necessary and sufficient for upregulation of NF-κB target genes early during HIV-1 infection in primary human MDMs. Jointly, our data support a model where both incoming and *de novo* HIV-1 Vpr induce DNA damage that activates ATM-NEMO signaling to upregulate NF-κB transcription (Fig. 7).

**Fig 7:**
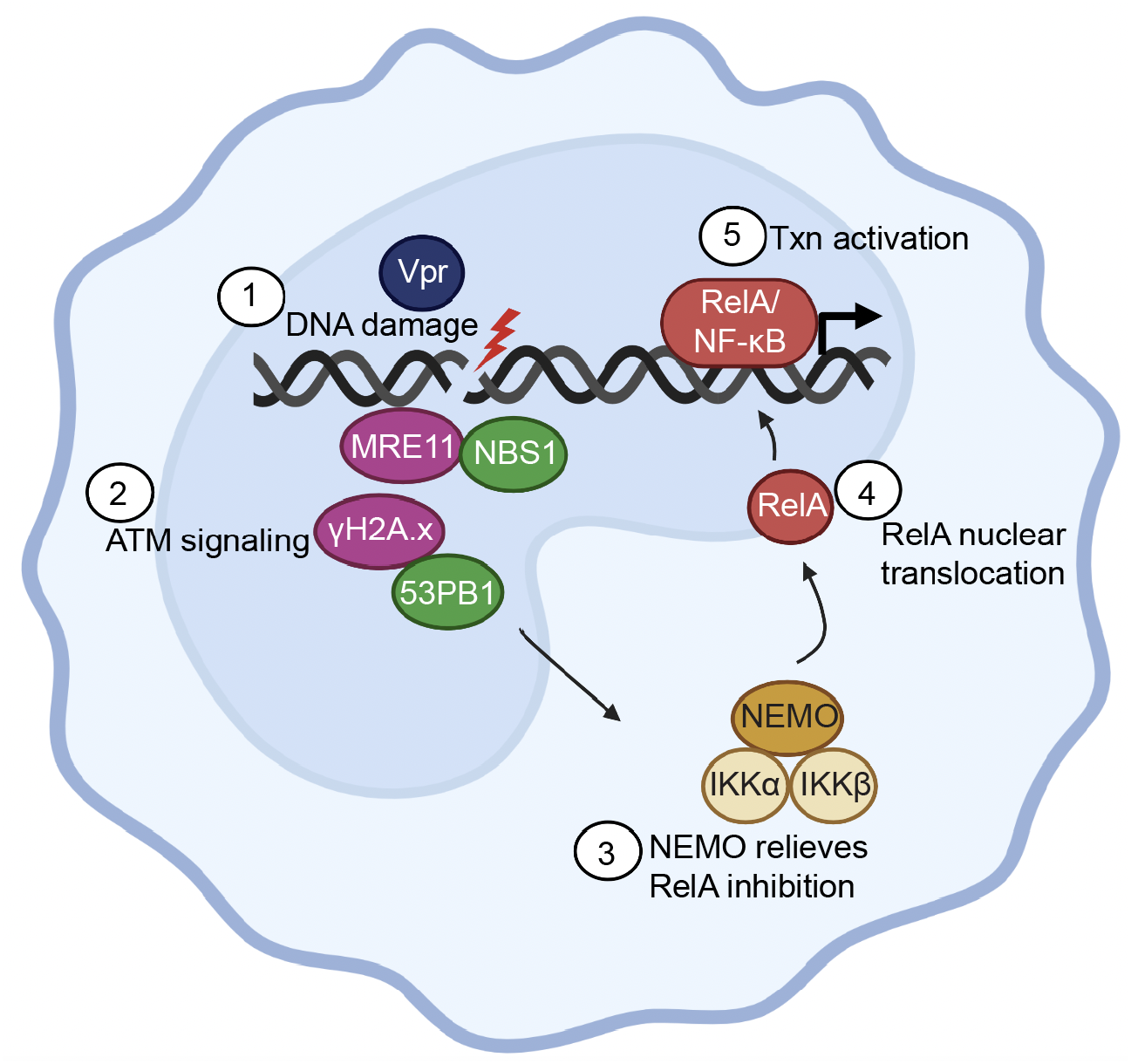
Model showing Vpr-induced DNA damage activates ATM and NEMO signaling resulting in RelA nuclear translocation and NF-κB transcriptional upregulation. 1. Vpr induces double-strand DNA breaks independent of cell cycle arrest. 2. Vpr-induced DNA damage activates markers of ATM signaling including MRE11, NBS1, γH2A.x, and 53PB1. 3. ATM-DDR activation results in NEMO relieving RelA inhibition from inhibitor of NF-κB (IκBα and IκBβ). 4. RelA translocates into the nucleus. 5. RelA binds to NF-κB promoters and activates transcription (txn) of NF-κB target genes such as BIRC3 and CXCL8. (Schematic made on Biorender).

## DISCUSSION

In this study, we tested the hypothesis that HIV-1 Vpr alters cellular transcription through induction of DNA damage and activation of DDR signaling. By leveraging Vpr mutants that separate Vpr-induced DNA damage from cell cycle arrest, we identified a current model where Vpr-induced DNA damage activates ATM-NEMO signaling, which stimulates RelA and upregulates NF-κB mediated transcription. Using U2OS and primary human MDMs, the latter of which allowed us to investigate how HIV-1 Vpr engages the DDR in this important cell type for the first time, we showed that Vpr induces DNA breaks and activates markers of ATM signaling independent of cell cycle arrest. Moreover, we showed that Vpr-induced DNA damage correlates with RelA/NF-κB activation, as assayed by bulk RNA-seq, and RelA immunofluorescence. We validated the upregulation of two NF-κB target genes, BIRC3 and CXCL8, and showed that inhibition of NEMO ablates the ability of Vpr to upregulate NF-κB transcription. Finally, VLP delivery of Vpr and HIV-1 infection show that incoming virion-associated Vpr is sufficient to induce DNA damage and activate RelA/NF-κB transcription. In complement, rAAV expression of Vpr show that *de novo* expressed Vpr can also induce DNA damage and activate RelA/NF-κB transcription. These data suggest that Vpr-induced DNA damage can play roles in both early and late stages of viral replication to alter cellular transcription. Overall, our data support a model where Vpr-induced DNA damage activates ATM-NEMO signaling and upregulates RelA/NF-κB transcription in cell lines and primary human MDMs.

Although Vpr is a multifunctional and enigmatic protein, the DNA damage response is central to many of the phenotypes associated with Vpr. Previous reports from our lab and others have worked to untangle how and why Vpr engages the DDR. A consensus is emerging that Vpr engages the DDR at multiple, potentially unique, steps. Through recruitment of the CRL4A^DCAF1^ ubiquitin ligase complex, Vpr degrades various DDR proteins, activates ATR signaling, represses double-strand DNA break repair, and causes cell cycle arrest (16,47). As shown here, Vpr-induced DNA damage directly correlates with NF-κB transcriptional activation that is independent of cell cycle arrest and dependent on ATM signaling. We have demonstrated that Vpr activates markers of ATM signaling, such as MRE11, NBS1, γH2A.x, and 53PB1, in a manner that resembles host activation rather than dysregulation. We have also shown that DNA damage and DDR activation occur in primary human MDMs and with virion-delivered, physiologically relevant levels of Vpr protein. Antagonism or dysregulation of DNA damage sensors through relocalization or sequestration is a conserved mechanism among many viruses to evade innate immune detection (48). While we have shown that Vpr-induced DNA damage activates markers of ATM signaling in a manner that seems to facilitate classical ATM activation, with the end goal of activating transcription, it remains to be seen whether Vpr-mediated activation of ATR, which correlates with CRL4A^DCAF1^ recruitment and cell cycle arrest, is classically activated or dysregulated. Moreover, how Vpr induces DNA breaks leading to both ATM and ATR activation is not understood.

Complementary studies have shown that Vpr modulates cellular (49) and viral transcription (50,51). In the context of cellular transcription, Vpr activates genes associated with innate immunity and proliferation in CD4+ T cells (52) and promotes the expression of proinflammatory cytokines in MDMs and monocyte-derived dendritic cells (MDDCs) (14,53). Together this suggests that Vpr has a vital role in transcriptional reprogramming to potentially create a proinflammatory environment that is conducive for viral replication. Consistent with these studies, we identified by bulk RNA-seq that Vpr regulates innate immune pathways that are either independent of or dependent on cell cycle arrest.

Our data also indicate that Vpr activates the RelA/NF-κB immune pathway without induction of cell cycle arrest. This is consistent with previous reports showing that Vpr activates NF-κB transcription via phosphorylation of TAK1 (24), an upstream regulator of NF-κB. We further found that Vpr-induced DNA damage upregulates RelA/NF-κB transcription via ATM-NEMO signaling, adding to the mechanistic understanding of NF-κB activation by Vpr. Through engagement with the CRL4A^DCAF1^ complex, Vpr has also been found to repress NF-κB activation by altering the availability of the NF-κB p50-p65 heterodimer, thus limiting proinflammatory cytokine expression (25). While it remains unclear what differentiates Vpr-mediated NF-κB activation from repression, whether Vpr engages the CRL4A^DCAF1^ complex may be a distinguishing factor. Furthermore, it is clear that in all cases Vpr carefully modulates NF-κB activation without globally activating interferon, which would inhibit viral replication. Thus, further studies understanding how Vpr manages to activate only aspects of NF-κB signaling (54,55) will help to define how Vpr may contribute to the ability of HIV to subvert the innate immune response.

Although various studies have shown that Vpr also alters viral transcription, the mechanisms are unclear and disparate. For example, Vpr-mediated LTR activation has been associated with cell cycle arrest, as the LTR is most transcriptionally active in G2/M (56,57), the degradation of host proteins such as CCDC137 (58), and CRL4A^DCAF1^ independent mechanisms (59). However, Vpr also promotes LTR transcription in noncycling cells (60,61) where induction of cell cycle arrest is absent and the necessity for degradation of specific host proteins in LTR activation has not been extensively examined. One benefit of Vpr activating NF-κB signaling via ATM is the potential direct enhancement of LTR transcription, as the HIV-1 LTR contains multiple NF-κB binding sites essential for viral gene expression. In addition to NF-κB, we further identified that Vpr activates CEBPB and JUN transcription factors, which have known roles in LTR activation. This potential direct effect on the LTR is similar to Vpr-mediated de-repression of the LTR by removal of the transcriptional repressor ZBTB2 following ATR activation (62). Overall, in the absence of cell cycle arrest, Vpr-induced DNA damage globally alters cellular transcription and upregulates NF-κB signaling and transcription factors that promote LTR-driven transcription, suggesting that Vpr-induced DNA damage is also important for promoting LTR transcription to enhance viral replication.

Together, our data suggest that during HIV infection incoming Vpr and *de novo* expressed Vpr prime the cellular environment by activating RelA/NF-κB signaling to promote transcription and enhance viral replication in macrophages. Our data align with the growing body of literature that supports the role of accessory proteins to modulate the host environment and the DDR to promote viral replication.

## MATERIALS AND METHODS

### Plasmids

pcDNA-3xFLAG-Vpr and pscAAV-mCherry-T2A-Vpr WT and mutant plasmids were generated as previously described (16). For rAAV production, pHelper and pAAV-2.5 capsid plasmids were used (Addgene and (16). For VLP production, psPAX2 and pmD2.G were used (Addgene). For HIV-1 ΔENV production, Bru-GFP ΔENV was generated as previously described (63) with pmD2.G (Addgene).

### Generation of Viruses

rAAV vectors packaging the pscAAV-mCherry-T2A-Vpr WT or mutant plasmids were generated by transient transfection of HEK 293 cells using polyethyleneimine (PEI) as previously described (64). Virus-like particles (VLPs) packaging Vpr WT or mutant proteins were generated by transient transfection of HEK 293T cells using TransIT-LT1 (Mirus). VLPs were harvested 48 hrs post transfection, concentrated through a 25% sucrose cushion at 24,000 rpm for 3 hrs at 4°C, and resuspended in PBS. Protein packaging was validated through western blot. HIV-1 ΔENV pseudotyped with VSV-G were generated by transient transfection of HEK 293T cells using TransIT-LT1 (Mirus). For HIV-1 ΔENV ΔVpr with Vpr WT packaged in trans (HIV-1ΔVpr +Vpr WT), 40 µg of pcDNA-3xFLAG-Vpr WT was also co-transfected in producer 293T cells. Virus was collected 48 hrs post media change and filtered through a 0.45 μm PES filter. Viral titer was identified by measuring activity of reverse-transcription through qPCR (65).

### Cell Lines and Cell Culture

Human bone osteosarcoma epithelial (U2OS), Human embryonic kidney (HEK) 293, and HEK 293T cells were cultured as adherent cells directly on tissue culture plastic (Greiner) in Dulbecco’s modified Eagle’s medium (DMEM) growth medium (high glucose, L-glutamine, no sodium pyruvate; Gibco) with 10% fetal bovine serum (FBS) (Gibco) and 1% penicillin-streptomycin (Gibco) at 37°C and 5% CO_2_. All cells were harvested using 0.05% trypsin-EDTA (Gibco). The panel of U2OS cells stably expressing 53BP1-GFP (43) and NBS1-GFP (42) were kindly provided by Claudia Lukas (University of Copenhagen, Denmark). The U2OS parental and NEMO knockout cells were kindly provided by Zhijian J. Chen (University of Texas Southwestern Medical Center). U2OS cells were pretreated with 50 µM of Nemo-Binding inhibitor (NBD) peptide or NBD negative control peptide (Thermo) for 2 hours prior to infection with rAAV or VLPs.

### Monocyte-Derived Macrophages (MDMs)

Human peripheral blood mononuclear cells (PBMCs) were obtained from human donors at the UCLA/CFAR Virology Core Laboratory. Primary monocytes were isolated from PBMCs by negative selection using the EasySep™ Monocyte Isolation Kit (STEM CELL) and were differentiated into monocyte-derived macrophages (MDMs) by stimulation with 20 ng/mL macrophage colony-stimulating factor (M-CSF) (R&D Systems). MDMs were cultured in Roswell Park Memorial Institute (RPMI) 1640 growth medium (L-Glutamine) with 10% fetal bovine serum (Gibco) at 37°C and 5% CO_2_ for 7 days while replenishing media every 3 days.

### Alkaline Comet Assay

The alkaline comet assay and data analysis was performed as previously described (16), with minor changes. MDMs were infected with VLPs delivering Vpr WT and mutants at equal protein levels or 25 µM Mitomycin C (Cayman). Cells were harvested with Accutase (STEM CELL) at 10 hrs post infection and resuspended in 0.5% low-melting-point agarose at 37°C. Images were acquired on the Zeiss Axio Imager Z1. Images were analyzed using the OpenComet plug-in for ImageJ.

### RNA-sequencing

Total RNA from U2OS cells was isolated using TRIzol® reagent (Invitrogen). RNA integrity was assessed using the Bioanalyzer TapeStation 4200 RNA High Sensitivity (Agilent). Library Preparation was completed using the KAPA mRNA HyperPrep Kit (Illumina) enriching for Poly(A) RNA with magnetic oligo-dT beads and attaching unique dual indexed adapters. Quality control of the library was done with the Bioanalyzer TapeStation 4200 D1000 High Sensitivity (Agilent). RNA was sequenced using the Hiseq3000 1×50 at the UCLA Technology Center for Genomics & Bioinformatics core. Reads were aligned to the Human h38 STAR genome and gene counts were determined using edgeR. log_2_ reads per kilobase of transcript per million reads mapped (rpkms) were calculated for samples compared to untreated cells. Heat map was generated using pheatmap hierarchical clustering on z-score log2 rpkms. Gene ontology, Protein-protein Interaction Enrichment Analysis, and TRRUST analysis were done on Metascape.

### Quantitative Reverse Transcription PCR (qRT-PCR)

Total RNA was isolated using the PureLink^TM^ RNA Mini Kit (Invitrogen). RNA was reverse transcribed using SuperScript IV First-Stand Synthesis System (Invitrogen) with Oligo(dT) primers. qRT-PCR was performed with PowerTrack SYBR Green Master Mix (Thermo Fisher Scientific) on the LightCycler 480 System (Roche) with the following primers (5’ to 3’) BIRC3 AAGCTACCTCTCAGCCTACTTT and CCACTGTTTTCTGTACCCGGA, CXCL8 TTTTGCCAAGGAGTGCTAAAGA and AACCCTCTGCACCCAGTTTTC, TNFα CTCTTCTGCCTGCTGCACTTTG and ATGGGCTACAGGCTTGTCACTC, HIV-1 unspliced RNA Genome GCGACGAAGACCTCCTCAG and GAGGTGGGTTGCTTTGATAGAGA, GAPDH CAAGATCATCAGCAATGCCT and AGGGATGATGTTCTGGAGAG, and Vpr AGGCCATGGCTTCATGGATTA and GATCTACCGGGTCCATTCCTG. mRNA levels were quantified by calculating ΔΔCt. Target transcript Ct values were normalized to the Ct value of the housekeeping gene GAPDH followed by calculating fold change to untreated or empty vector treated cells.

### Live-Cell Imaging

U2OS-53PB1-GFP and U2OS-NBS1-GFP cells were imaged using the IncuCyte S3 Live-Cell Analysis Instrument (Sartorius). Mean Fluorescence Intensity (MFI) was calculated using the Sartorius software. For higher resolution, U2OS-53PB1-GFP and U2OS-NBS1-GFP cells were imaged using the LSM 900. Foci per cell and foci size were analyzed using ImageJ.

### Immunofluorescence

Cells were plated in 6- or 24-well tissue culture plates (Greiner) and allowed to adhere overnight, then infected with VLP (equal protein expression), rAAV-2.5 (1.4 × 10^8^ copies/well), or etoposide (Sigma). U2OS cells were permeabilized with 0.5% Triton X-100 in PBS at 4°C for 5 min and fixed in 4% PFA for 20 min, MDM cells were permeabilized with 0.1% Saponin (Fisher) in PBS at 4°C for 15 min and fixed in 4% PFA for 15 min. Cells were washed, incubated with blocking buffer (3% BSA, 0.05% Tween 20, and 0.04 NaN_3_ in PBS for U2OS cells or 3% FBS in 0.1% Saponin for MDM cells) for 30 min. Cells were probed with appropriate primary antibodies (anti-γH2A.x Ser139, anti-53BP1, or anti-RelA(p65) [Cell Signaling], anti-GFP [Takara], and anti-MRE11 [Novous]) and then washed and probed with Alexa Fluor-conjugated secondary antibodies (Life Technologies). Nuclei were stained with diamidino-2-phenylindole (DAPI; Life Technologies). Images were acquired on the LSM 980.

### Western Blots and Co-immunoprecipitations

Protein was collected from cells as previously described (16). For co-immunoprecipitations, cells were washed with cold phosphate-buffered saline (PBS) and lysed with radioimmunoprecipitation assay (RIPA) buffer (50 mM Tris-HCl [pH 8.0], 200 mM NaCl, 1 mM EDTA, 0.1% SDS, 1% NP-40, 0.5% sodium deoxycholate, Benzonase, protease inhibitor) and clarified by centrifugation 15,500 × g for 15 min. Immunoprecipitations were performed overnight at 4°C with Dynabeads Protein G beads (Invitrogen) conjugated to mouse anti-FLAG M2 (Sigma-Aldrich). For western blotting, samples were boiled in 4× sample buffer (40% glycerol, 240 mM Tris, pH 6.8, 8% SDS, 0.5% β-mercaptoethanol, and bromophenol blue) in preparation for SDS-PAGE using 12% Bis-Tris polyacrylamide gels and subsequently transferred onto a polyvinylidene difluoride membrane. Membranes were blocked in intercept blocking buffer (Li-COR Biosciences). Immunoblotting was performed using mouse anti-FLAG M2 (Sigma-Aldrich), rabbit anti-Actin (Bethyl), mouse alpha-Tubulin (Sigma), or rabbit anti-DCAF1 (Proteintech) for 1 hr. Blots were incubated with secondary antibodies IRDye 800CW anti-Rabbit and IRDye 680RD anti-Mouse (Li-COR Biosciences) for 1 hr and then visualized using the Li-COR Odyssey M (Li-COR Biosciences).

### HIV-1 ΔEnv Infection

MDMs were plated in 96-well tissue culture plates (Greiner) and allowed to adhere overnight. MDMs underwent spinfection with HIV-1 ΔEnv pseudotyped with VSV-G (HIV-1), HIV-1 ΔEnv ΔVpr pseudotyped with VSV-G (HIV-1 ΔVpr), HIV-1 ΔEnv pseudotyped with VSV-G packaging Vpr WT (HIV-1 ΔVpr +Vpr WT) or mock at 1200× *g* for 90 min at 37°C. Infection was assessed 48 hours after infection via qRT-PCR and Flow Cytometry.

### Flow Cytometry

Isolated monocytes and infected MDMs were lifted from tissue culture plates using accutase (STEM CELL). Cells were stained for CD14-FITC, CD45-APC, or CD16-APC (STEM CELL) for 30 min at 4°C, washed with PBS, fixed in 4% PFA for 15 min, permeabilized with 0.1% Triton X-100 in PBS at 4°C for 15 min, then washed with PBS. Cells were probed for HIV-1 core antigen-FITC KC57 (Beckman Coulter) for 1 hr at 4°C then washed with PBS and resuspended in FACS buffer (5% FBS in PBS). Events were assessed by flow cytometry on the AttuneNxT (Thermo Fisher Scientific). At least 10,000 events were collected per run. Data was analyzed using FlowJo software.

### Statistical Analyses

All statistical analyses were performed using GraphPad Prism 9.

## Data access

The bulk RNA-sequencing data in this publication was deposited in NCBI’s Gene Expression Omnibus and can be accessed through GEO series accession number GSE253779 (https://www.ncbi.nlm.nih.gov/geo/query/acc.cgi?acc=GSE253779).

## AUTHOR CONTRIBUTIONS

Conceptualization, C.S. and O.I.F; methodology, C.S. and K.N.; writing original draft C.S and O.I.F; writing, review and editing C.S., K.N., and O.I.F; visualization C.S. and O.I.F.; supervision and funding acquisition O.I.F. All authors have read and agreed to the published version of the manuscript.

## ACKNOWLEDGEMENTS

We like to thank Dr. Julia Mack and Julianne Ashby for their assistance and expertise with Live-cell imaging using the LSM900. We thank Dr. Alexander Hoffmann and Dr. Diane Lefaudeux for instruction on RNA-seq analysis. We thank Dr. Claudia Lukas for providing U2OS cells that stably express 53PB1-GFP and NBS1-GFP. We thank Dr. Zhijian J. Chen for providing U2OS NEMO knockout cells. We thank Dr. Steven Bensinger for the LightCycler 480 System, Dr. Jesse Zamudio for the Qubit fluorometer, Dr. Peter Bradley for the Zeiss Axio Imager Z1, Dr. Matteo Pellegrini for the TapeStation, Dr. Steve Jacobsen and Dr. Jefferey Long for the LSM980. We thank Dr. Randilea Nichols Doyle and Vivian Yang for providing comments on the manuscript.

## SUPPLEMENTAL FIGURE LEGENDS

**Fig S1:**
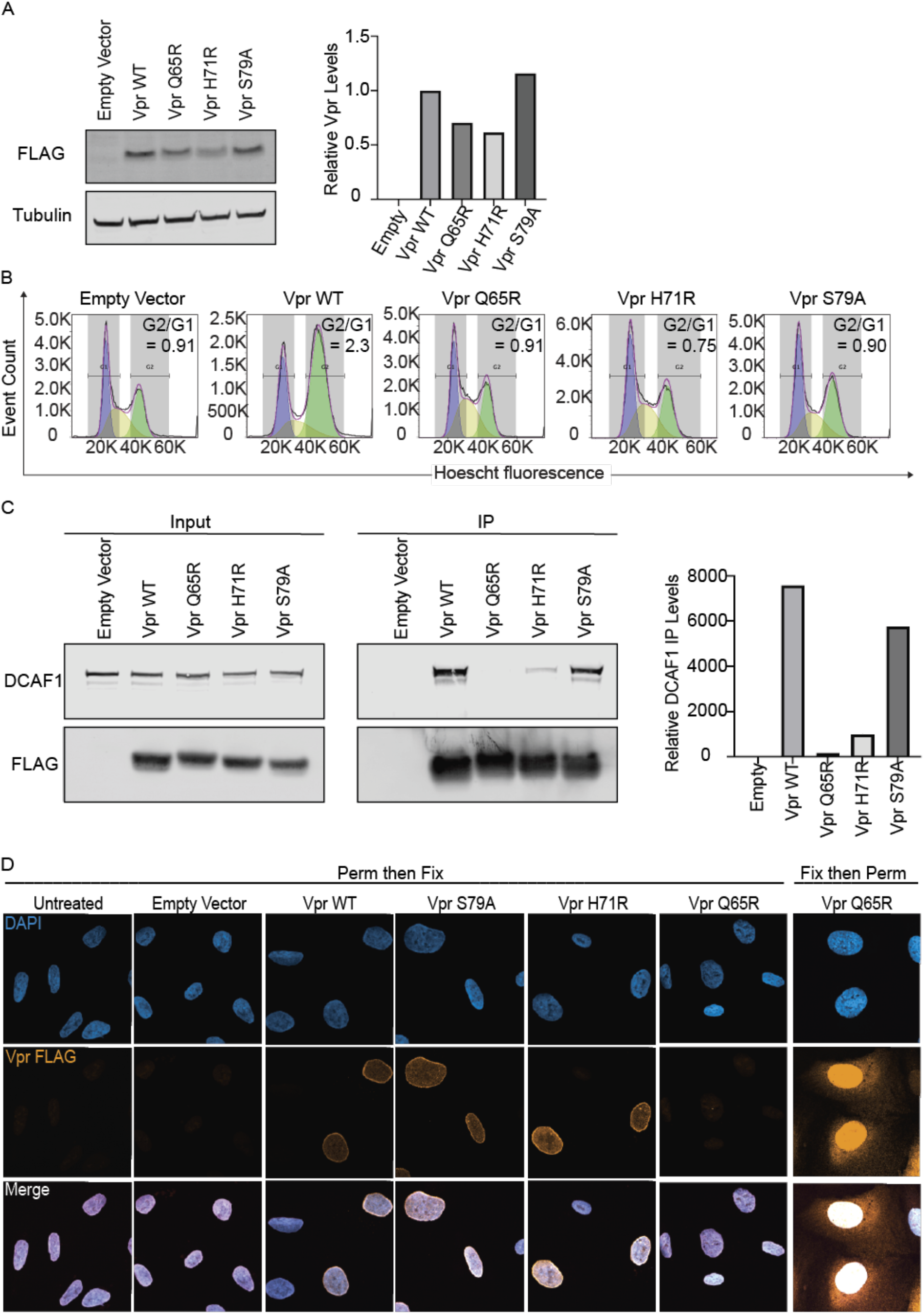
Characterization of HIV-1 Vpr mutants S79A, H71R, and Q65R. (A) Representative western blot of 3X FLAG-Vpr WT and mutants from U2OS cells infected with rAAV expressing Vpr WT, Q65R, H71R, S79A, or empty vector (EV) (negative control) as in Fig. 1A. α-tubulin (loading control). Quantification of Vpr levels relative to corresponding α-tubulin and normalized to Vpr WT. (B) Cell cycle arrest analysis of U2OS cell treated as in Fig. 1A. Percent of events in G2 and G1 were measured using flow cytometry gating on mCherry positive cells (rAAV infection) and stained for Hoescht (DNA content). N=3, one representative experiment shown. (C) Co-immunoprecipitation against 3X FLAG-Vpr WT and mutants, probed for endogenous DCAF1, from U2OS cells infected under the same conditions as Fig. 1A. Quantification of DCAF1 immunoprecipitated with Vpr WT and mutants relative to light chain. N=3, one representative experiment shown. (D) Representative immunofluorescence images of U2OS cells infected under the same conditions as Fig. 1A. Cells were either permeabilized then fixed or fixed and then permeabilized. Vpr FLAG (orange) and DAPI (blue). Representative images taken at 24 hpi at 63x magnification. N=3, one representative experiment shown. Related to Figures 1-5.

**Fig S2:**
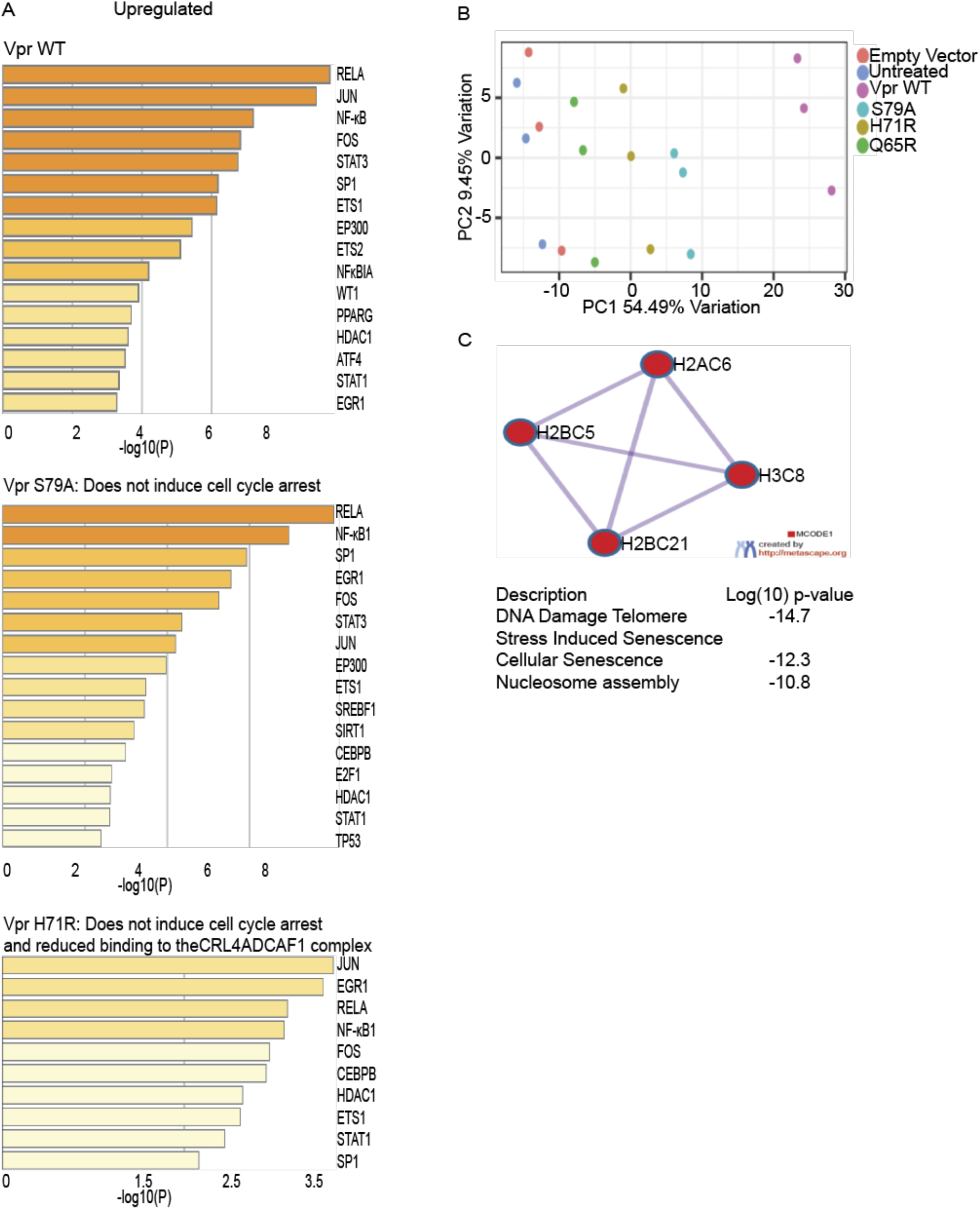
Vpr alters cellular transcription through diverse mechanisms that are either dependent or independent of cell cycle arrest. (A) Gene ontology using Metascape displaying TRRUST analysis for upregulated genes. (B) Principal component analysis (PCA) of the RNA-seq data showing the relatedness of the three biological replicates compared to empty vector or untreated cells. (C) Protein-protein Interaction Enrichment Analysis using Metascape of the shared downregulated genes among Vpr WT, S79A and H71R. Related to Figure 1.

**Fig S3:**
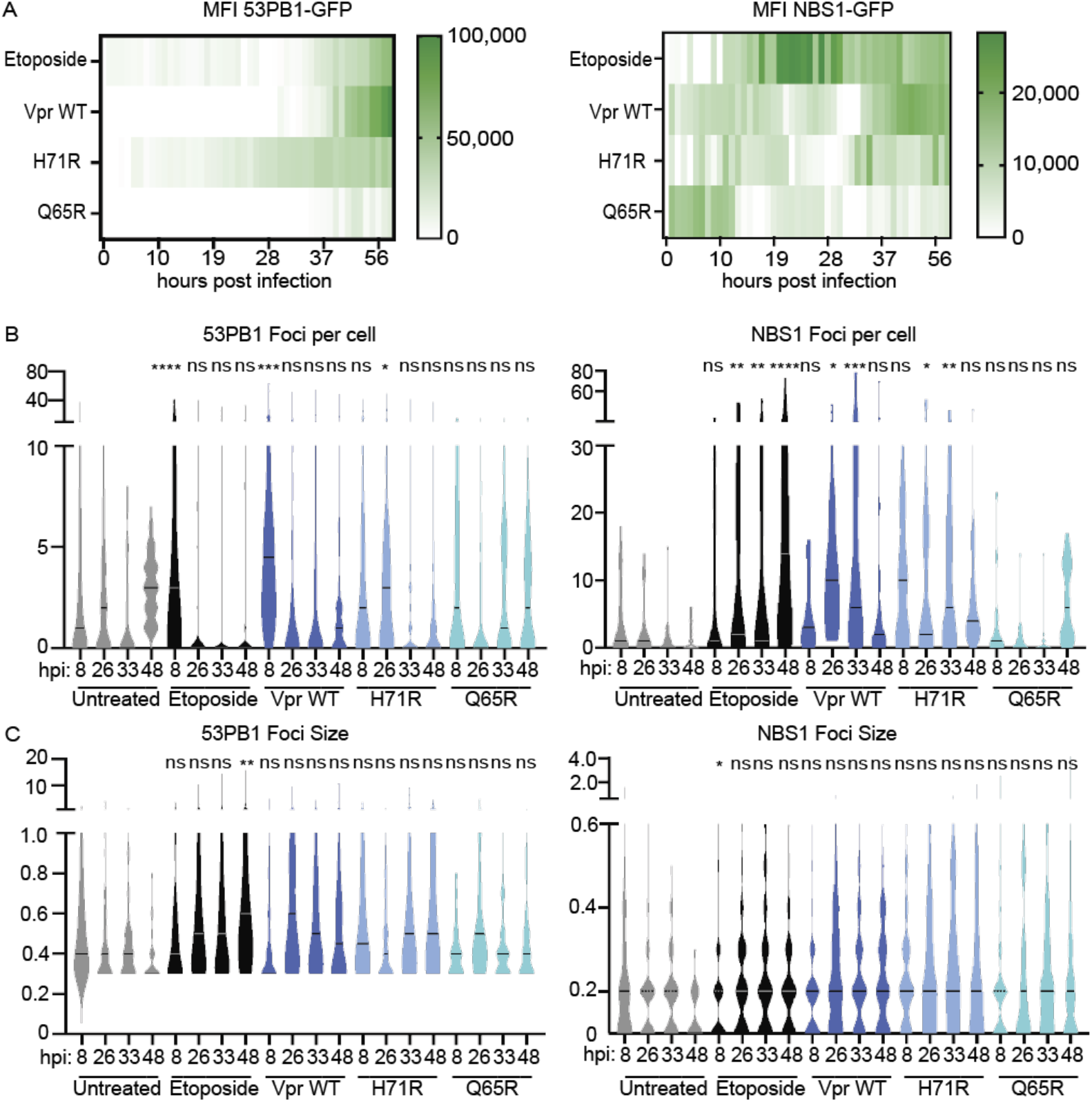
Vpr-induced DNA damage activates markers of ATM signaling. (A) Live-cell imaging taken on the IncuCyte of U2OS NBS1-GFP or 53PB1-GFP cells infected as in Fig. 1A for 56 hrs. Heatmap displays NBS1 and 53PB1 mean fluorescence activity (MFI) (green) as arbritary units (AU). N=2, one representative experiment shown. (B & C) Quantification of DNA damage induced in the live-cell imaging experiment in Fig. 3A at 8, 26, 33, and 48hpi. Foci per cell and foci size were quantified using ImageJ software. Images were taken at 63x magnification using the LSM 900. N=3, one representative experiment shown with at least 50 cells per condition. Arbitrary units (AU) displayed. Asterisk indicate statistical significance compared to untreated cells (negative control) at the corresponding timepoint, as determined by one-way ANOVA test (NS, nonsignificant; * P<0.03, ** P < 0.005, *** P< 0.0003, **** P< 0.0001). Related to Figure 3.

**Fig S4:**
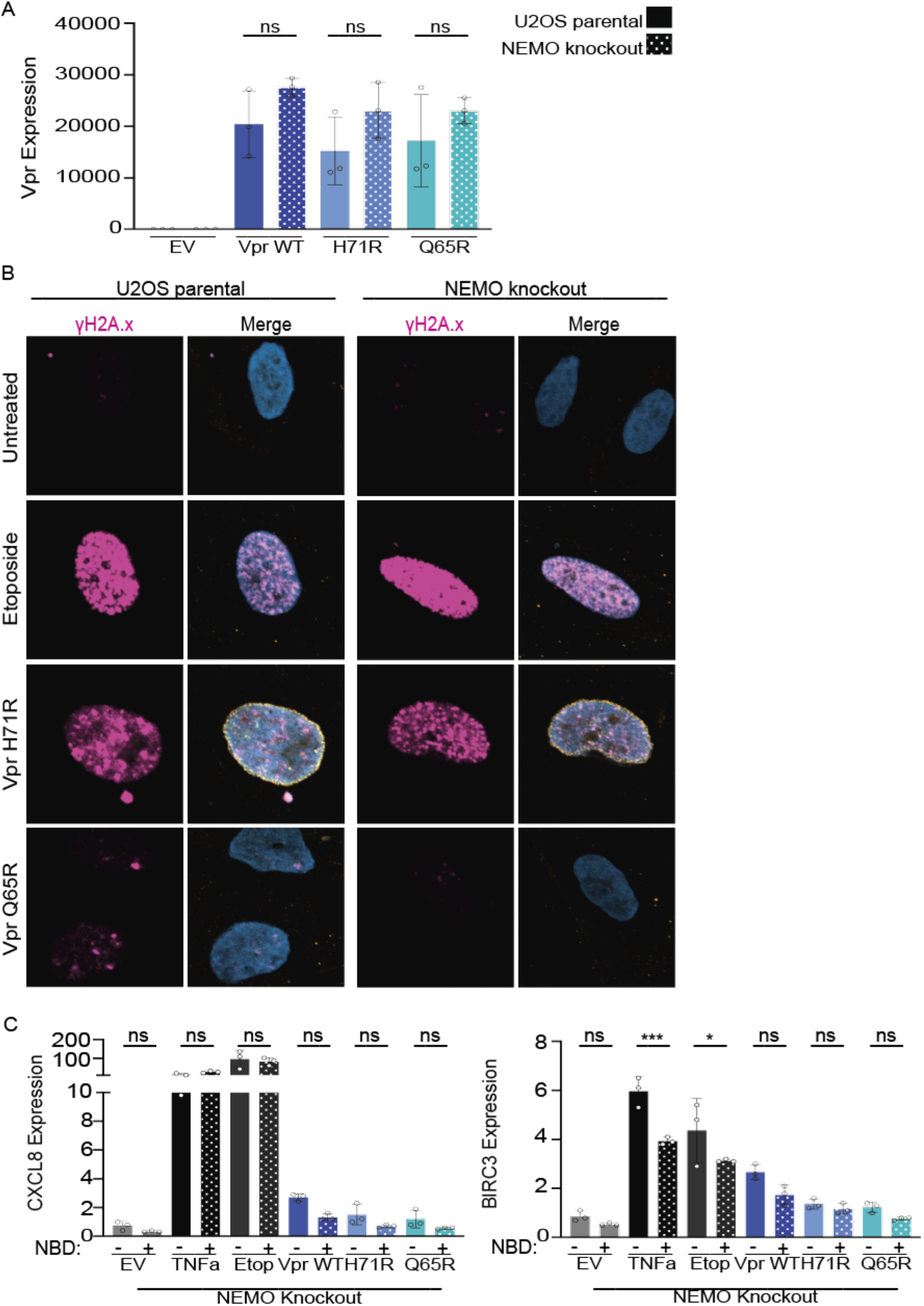
Nemo knockout and NEMO-Binding inhibitor peptide (NBD) comparably inhibit Vpr-induced NF-kB activation. (A) qRT-PCR for Vpr expression of U2OS cells infected as in Fig. 4 in U2OS parental or NEMO knockout cells with technical triplicates. (B) Representative images of U2OS parental or NEMO knockout cells infected as in Fig. 1A displaying γH2A.x (magenta), 3X FLAG-Vpr (yellow), DAPI (blue). Representative images taken at 24hpi at 63x magnification. N=3, one representative experiment shown. (C) qRT-PCR of BIRC3 and CXCL8 in NEMO knockout cells in the presence of Nemo-Binding inhibitor peptide (NBD, +) or the NBD negative control peptide (-) at 36 hpi. Cells were treated under the same conditions as Fig. 4. Normalized expression to GAPDH. N=3, one representative experiment shown. Asterisk indicate statistical significance compared to NBD negative control, as determined by one-way ANOVA test (NS, nonsignificant; * P< 0.003, *** P< 0.0003). Related to Figure 4.

**Fig S5:**
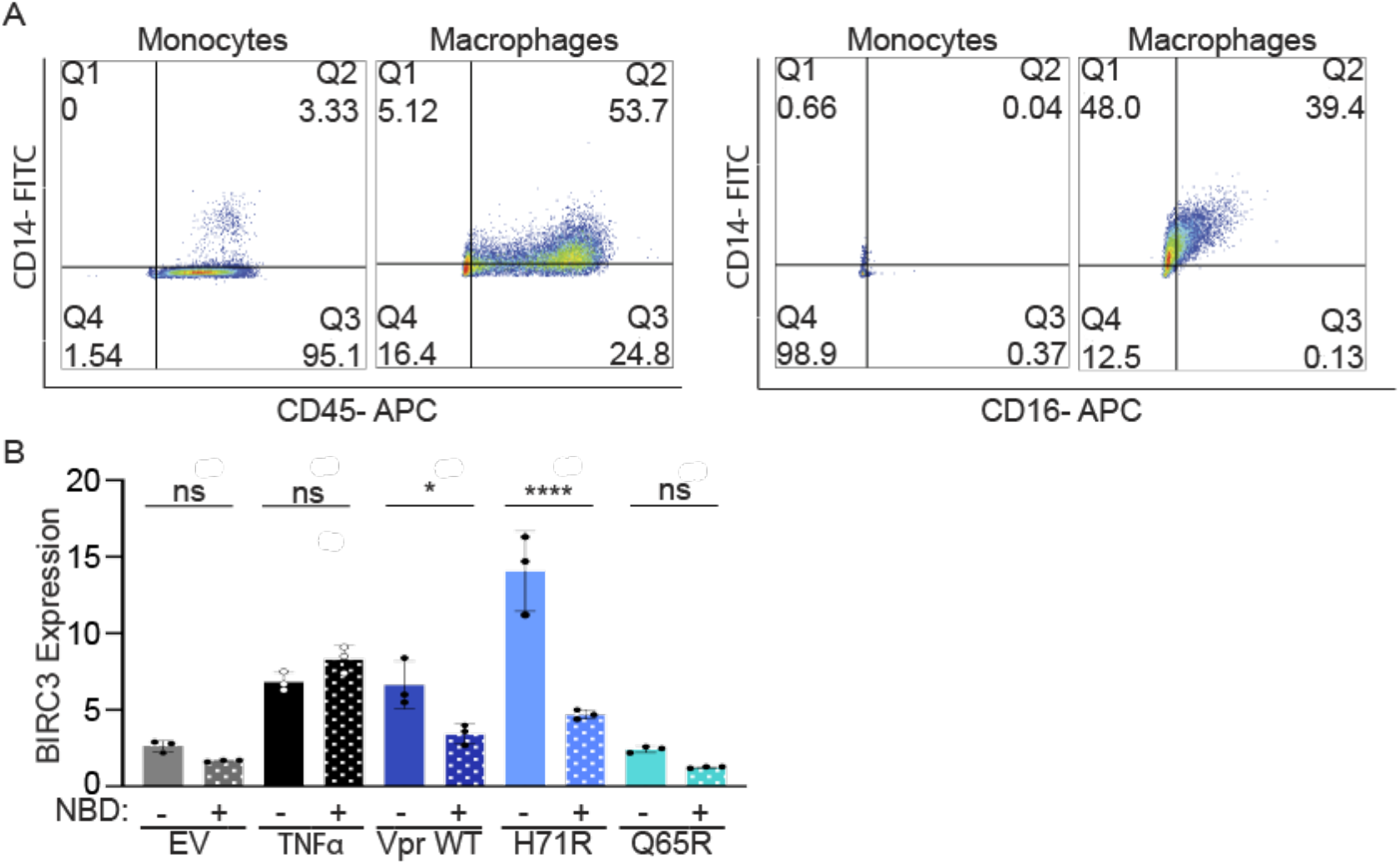
Infection of primary MDMs with VLPs packaging Vpr protein upregulates NF-κB transcription dependent on NEMO. (A) Flow cytometry plots displaying differentiation of PBMC-derived monocytes to macrophages. Primary MDMs are CD14+, CD45+ and CD16+. N=4 one representative experiment shown. (B) qRT-PCR for BIRC3 of MDMs treated with NBD inhibitor peptide as in Fig. 5C with TNFα as positive control. N=3, displaying mean of 3 separate donors. Asterisk indicate statistical significance compared to NBD negative control, as determined by one-way ANOVA test (NS, nonsignificant; * P< 0.003, **** P< 0.0001). Related to Figure 5.

**Fig S6:**
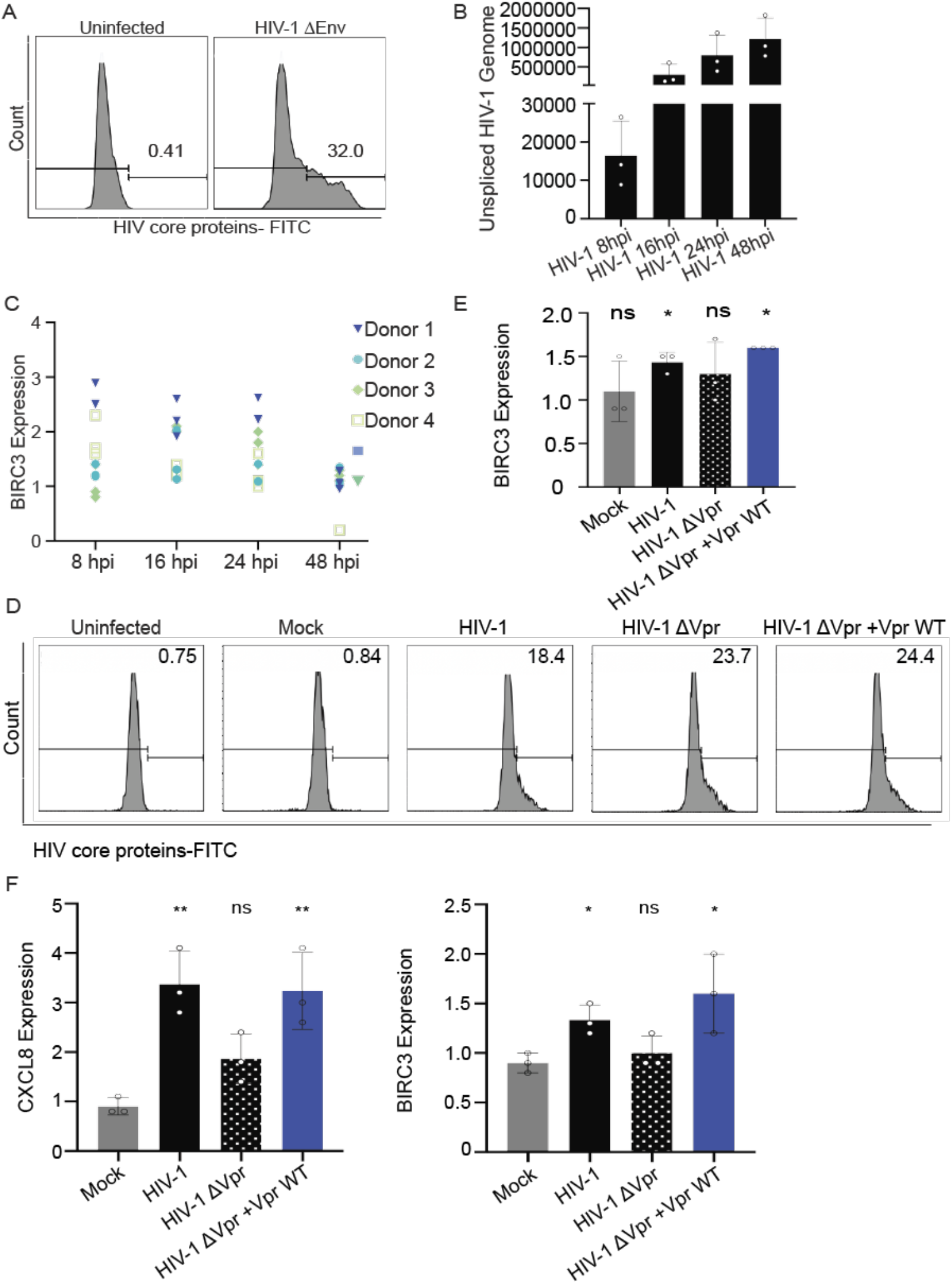
HIV-1 infection of primary human MDMs upregulates NF-κB transcription and is rescued by Vpr. (A) Histogram quantifying percent of infected MDMs with HIV-1 ΔEnv at 48hpi. N=4, one representative experiment shown. (B) qRT-PCR for unspliced HIV-1 RNA at 8, 16, 24 and 48hpi. N=4, one representative experiment shown. (C) qRT-PCR for BIRC3 or CXCL8 of MDMs infected with HIV-1 ΔEnv at 8, 16, 24 and 48hpi in four separate donors. N=4. (D) Histogram quantifying percent of infected MDMs with HIV-1, HIV-1ΔVpr, HIV-1ΔVpr +Vpr, mock or untreated at 48hpi. N=3, one representative experiment shown. (E) qRT-PCR for BIRC3 of MDMs infected with MOI 5U/mL of HIV-1, HIV-1 ΔVpr, HIV-1 ΔVpr +Vpr WT viruses or mock. Displaying the mean of three separate donors. Normalized expression to GAPDH. Asterisk indicate statistical significance compared to untreated negative control, as determined by Welch t-test for BIRC3 (NS, nonsignificant; * P<0.04). (F) qRT-PCR for BIRC3 or CXCL8 of MDMs infected with HIV-1, HIV-1ΔVpr, HIV-1ΔVpr +Vpr, mock at 8hpi from three separate donors. One representative experiment shown with technical triplicates. Asterisk indicate statistical significance compared to untreated control, as determined by one-way ANOVA test (NS, nonsignificant; * P< 0.003, ** P< 0.001). Related to Figure 6.

**Files S1 & S2: HIV-1 Vpr WT and mutants alter cellular transcription.** Analysis of all upregulated (File S1) and downregulated (File S2) genes in the RNA seq analysis that passes statistical significance of Log2 (1.25), p < 0.01, and FDR < 4.50E-05. Counts per million reads mapped (CPM) Log2 fold changes were calculated relative to untreated control. Related to Figure 1 and Figure S2.

